# How the Brain Predicts Timing: Distinct Network Hubs for Predicting and Evaluating Auditory Sensory Events

**DOI:** 10.1101/2024.11.19.624242

**Authors:** Péter Nagy, Petra Kovács, Ádám Boncz, Orsolya Szalárdy, Robert Baumgartner, Karolina Ignatiadis, István Winkler, Brigitta Tóth

## Abstract

Temporal prediction enhances perceptual processing by aligning neural excitability with expected sensory events. While local oscillatory mechanisms are known to support timing, less is understood about how large-scale functional brain networks dynamically coordinate predictive processes. In particular, it remains unclear how functional connectivity (FC)—the integration of information into network hubs—differs during expectation formation (post-cue) versus outcome evaluation (post-target), and how this varies across levels of predictability. To investigate this, we recorded electroencephalogram (EEG) while participants performed a cued auditory target-detection task with varying temporal predictability (80% and 50%). Event-related potential (ERP) results revealed that implicit temporal predictability primarily modulated later evaluative processes (P3b, frontal negativity), rather than early sensory components, consistent with context updating under uncertainty. FC was analyzed using a data-driven approach based on Normalized Directed Transfer Entropy (NDTE) applied to EEG difference waveforms between high- and low-predictability conditions. Connectivity was examined separately for the post-cue and post-target periods to distinguish prediction and evaluation phases. Behaviorally, higher temporal predictability facilitated faster reaction times. Connectivity analyses revealed largely overlapping but somewhat distinct network dynamics for prediction and evaluation phases of the signal processing.

## 1 Introduction

To navigate the world efficiently, the brain must not only recognize what is happening, but also anticipate when it will happen. Temporal prediction—the ability to forecast the timing of sensory events—enables the nervous system to align moments of heightened neural excitability with expected stimuli, optimizing perception and behavior (Jones, 2018; Large & Jones, 1999; Nobre & van Ede, 2018). When the timing of stimuli is predictable, behavioral responses improve (e.g., faster reaction times, increased accuracy; Toosi et al., 2017), suggesting that predictive timing modulates neural dynamics to prioritize the relevant sensory input. Although local cortical oscillations in delta/theta bands support the processes of temporal prediction (Schroeder & Lakatos, 2009; Arnal & Giraud, 2012; Benedetto et al., 2021), less is known about how timing signals are communicated within large-scale brain networks. The present study addresses this gap by investigating how large-scale networks of cortical hubs dynamically coordinate during different phases of temporal prediction.

A growing body of research suggests that the mechanisms underlying temporal prediction involve both local oscillatory dynamics and interactions across large-scale brain networks. Cortical oscillations—particularly in the delta (0.5-4 Hz) and theta (4-8 Hz) bands—regulate temporal expectations by modulating excitability (Schroeder & Lakatos, 2009; Arnal & Giraud, 2012; Benedetto et al., 2021). For instance, the phase of low-frequency oscillations resets before an anticipated stimulus, and stronger phase alignment has been observed for temporally predictable events (Lakatos et al., 2008; Stefanics et al., 2010). These local entrainment processes are foundational for timing, but they do not fully explain how timing-related information propagates and is integrated across the brain.

Temporal prediction also engages large-scale functional connectivity (FC) networks that coordinate sensory and higher-order cortical regions (Arnal et al., 2015; Darriba & Waszak, 2018). Defined as the synchronization or statistical dependence of activity between distant brain regions, FC supports the flow of predictive information, possibly enabling top-down modulation from higher-order to sensory areas (Sauseng & Klimesch, 2008). While previous research has implicated both frontal and sensory cortices in temporal prediction (Coull et al., 2016; Meirhaeghe et al., 2021; de Lange et al., 2018), the specific architecture and directional roles of network hubs—especially their dynamic involvement during different temporal prediction phases—remain unresolved. Whether predictive timing signals originate predominantly in frontal hubs or emerge within sensory regions (Rao & Ballard, 1999; Kok et al., 2012; Alamia & VanRullen, 2019), or whether they are distributed across both (Hohwy, 2013; van Moorselaar & Slagter, 2019) is unclear. To trace these dynamics, we need methods that resolve when and where information flows occur.

A parallel line of research, reviewed by Korka et al. (2022), demonstrates that intention-based prediction has a strong influence on auditory processing—even in passive situations and for stimulus-omission paradigms—supporting the Auditory Event Representation System (AERS; Winkler & Schröger, 2015) theory. Korka and colleagues’ (2022) extended AERS model illustrates how action intentions elicit predictive signals transmitted from the frontal to the auditory cortices, operating across pre-attentive (e.g., MMN) to later evaluative stages. Critically, intention-generated predictions can override sensory regularities, emphasizing top-down influences on hierarchical processing. These findings raise the question: do intention- and stimulus-based predictions rely on common or distinct network hubs, and are they active before or after target onset?

There is moreover an ongoing theoretical debate regarding whether temporal predictions originate in higher-order frontal regions and are relayed to sensory cortices (e.g., predictive coding theory; Friston, 2005; Clark, 2013), or whether sensory areas can independently encode and utilize temporal regularities (Rao & Ballard, 1999; Kok et al., 2012; Alamia & VanRullen, 2019). Hybrid models propose a division of labor, with frontal areas providing top-down predictions and sensory cortices generating prediction errors or updating local representations (Hohwy, 2013; Nobre & van Ede, 2018; van Moorselaar & Slagter, 2019). Testing these competing models requires tools capable of tracking fast, distributed interactions across the cortex with high temporal precision.

In the current study, we aim to address these questions by characterizing the FC of cortical hubs during two distinct phases of temporal prediction: (1) the post-cue prediction phase, when participants form expectations about upcoming events, and (2) the post-target evaluation phase, when sensory evidence confirms or violates those expectations. This distinction allows us to investigate how predictive timing and feedback mechanisms evolve over time and whether the same or different network hubs are engaged at each stage.

To achieve this, we used electroencephalogram (EEG) and a cued auditory detection task with manipulated temporal predictability. Participants heard a cue that probabilistically predicted the timing (early or late) of an upcoming target sound. Based on this design, we defined two levels of temporal predictability: in the predictable condition, early targets followed the cue with high probability (80%), whereas in the non-predictable condition, the cue provided no reliable timing information (50%). We hypothesized that early targets preceded by predictive cues (80% predictable) would elicit faster reaction times (RTs) than early targets preceded by unpredictive cues (50% predictable), due to increased temporal predictability. By computing the difference waveforms between 50% and 80% predictable early targets and between the corresponding cues, we isolated activity associated with probabilistic timing. FC analyses were conducted on these difference-signals, separately for the post-cue and post-target periods.

Here, we apply a hub-based connectivity approach to capture the directional flow of predictive information across the cortex with high temporal precision (Deco et al., 2021; Ignatiadis et al., 2024). Hubs that are more densely functionally connected among themselves than to other brain regions, from which they receive integrative information and whose outflow to different areas is sparse, comprise core sets that constitute a functional rich club (FRIC). This approach complements traditional node-based or symmetric FC methods by providing a directional view of network dynamics (Fiebelkorn et al., 2019; Ignatiadis et al., 2024). The FRIC approach allows us to contrast two theoretically opposing hypotheses: 1) The frontal-dominant hypothesis: Predictive timing relies primarily on inflow to frontal hubs post-cue (formation of expectations) and outflow to sensory cortices post-target (error resolution), consistent with hierarchical predictive coding (Friston, 2005; Alamia & VanRullen, 2019). 2) The distributed integration hypothesis: Both frontal and sensory hubs receive predictive input across phases, with inflow distributed more symmetrically across the cortex, consistent with local prediction models (Kok et al., 2012; Benedetto et al., 2021). By contrasting post-cue and post-target FC topographies, we also assessed whether temporal predictions rely on the same hub architecture across time or whether the prediction network reorganizes dynamically during expectation formation vs. sensory updating.

In summary, this study presents a novel investigation into the temporal evolution of predictive networks by applying FRIC-based FC analysis to EEG. Our goal is to clarify the directionality and dynamics of cortical involvement in perceptual prediction, shedding light on how distributed brain regions flexibly coordinate to support temporal anticipation.

## 2 Methods

### 2.1 Participants

Twenty healthy young adults participated in the study, recruited from a pool of participants who had previously participated in our experiments (10 female; mean age: 22.4 ± 2.35 years, all right-handed). Participants were financially compensated for their participation (approx. 4 euros per hour). None of the participants reported any neurological diseases or hearing problems. Participants signed written informed consent forms after the aims and procedures of the experiment were explained. The study adhered to the Declaration of Helsinki guidelines and was approved by the local ethics committee of the Institute of Cognitive Neuroscience and Psychology at the HUN-REN Research Centre for Natural Science (United Ethical Review Committee for Research in Psychology; EPKEB). The EEG data of 3 participants were excluded from the analysis due to technical issues (missing blocks) during data acquisition. The final group for which EEG analysis was conducted consisted of 17 listeners (10 female and 7 male), with a mean age of 22.6 ± 2.45 years.

### 2.2 Stimuli and procedure

Participants performed an auditory target detection task while EEG was recorded. They were instructed to detect target tones as quickly as possible by pressing a key on a computer keyboard with the index finger of their dominant hand (all participants were right-handed). Each target was preceded by an auditory cue (Figure 1A).

**Figure 1.**
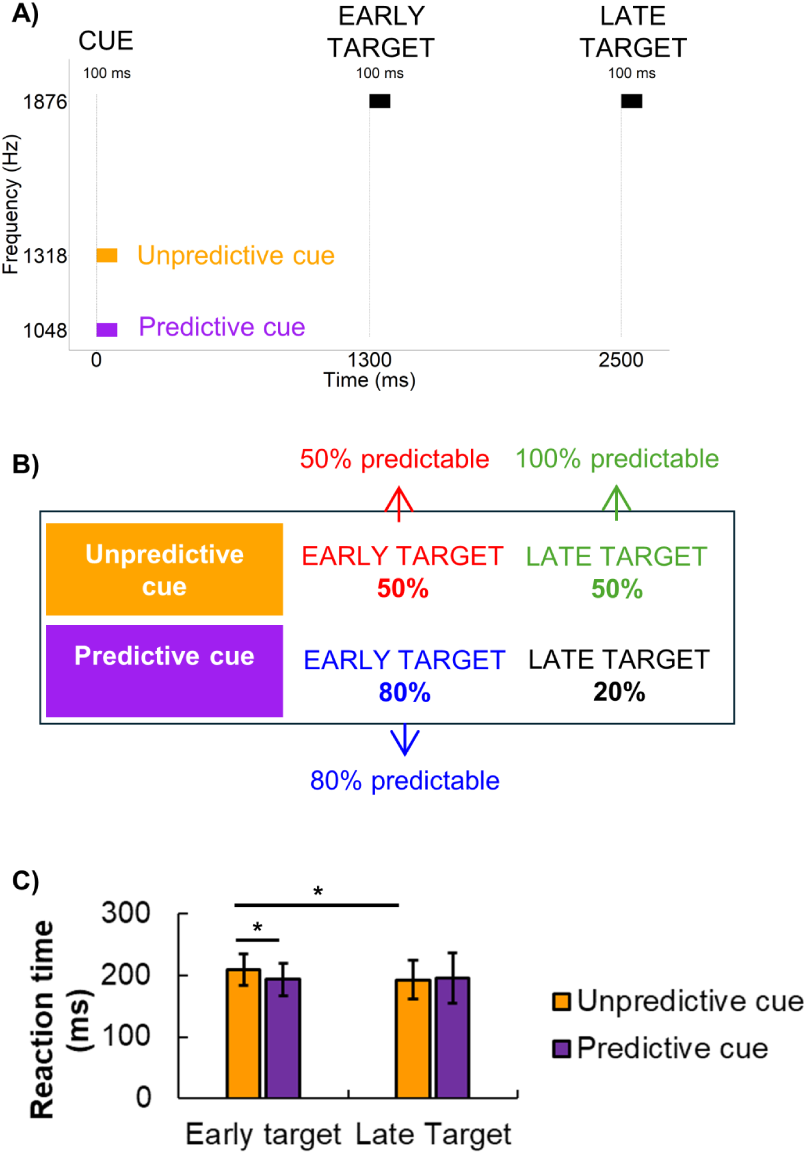
Stimuli, conditions, and reaction time results. A) Timeline of the stimulus presentation. In each trial, one cue and one target were presented, each lasting 100 ms. The target followed the cue after either a short or a long ISI (early vs. late target). Two types of cues were used: unpredictive and predictive cues. The cues differed in their pitch. B) After a predictive cue, an early target followed with 80% certainty, and after an unpredictive cue, early and late targets followed with equal likelihood. We refer to the 50%, 80%, and 100% predictable conditions, based on their within-trial probability, as shown in the figure. C) Mean RTs to the target events and the results of the comparisons between them (horizontal lines with an asterisk above them denote significant differences; error bars denote standard deviation).

The interval between the cue and the target—referred to as the inter-stimulus interval (ISI)— was either 1200 ms or 2400 ms, corresponding to *early* and *late* target conditions, respectively. The inter-trial interval (ITI) varied randomly between 200 and 1000 ms to minimize temporal predictability across trials. All stimuli were presented at a uniform sound pressure level of approximately 70 dB.

Two distinct cue tones were used to manipulate the *predictability* of the target’s timing. These tones had frequencies of 1048 Hz and 1318 Hz, each presented for 100 ms with 10 ms linear rise and fall times. The target tone was always 1876 Hz, with a duration of 100 ms, including 10 ms rise and fall times. The cue frequencies were separated by one semitone, while the target tone was two semitones above the higher cue, ensuring clear perceptual distinction.

One cue served as a predictive cue, while the other served as a non-predictive cue, with the assignment counterbalanced across participants. After the predictive cue, the target was followed at a short ISI (1200 ms) on 80% of the trials, making early targets more likely than late ones. In contrast, the non-predictive cue was followed by early and late targets with equal probability (50%). Participants were not informed whether a cue was predictive. Based on this design, we identified three levels of temporal predictability for target events: **1) 50% predictable**: Early targets following a non-predictive cue. **2) 80% predictable**: Early targets following a predictive cue. **3) 100% predictable**: Late targets, which occurred with certainty (100%) if an early target did not appear after the cue. Each participant completed three blocks of 50 trials per cue type, resulting in a total of 300 target trials (150 per cue type). The condition order was randomized across participants. Before the main experiment, participants completed a brief familiarization session with the stimuli and task.

This classification allowed us to examine how predictability modulates neural and behavioral responses (Figure 1A). However, since the 100% condition involved a longer delay before the target, it was not directly comparable to the early target conditions, i.e., due to response selection, preparatory processes, and movement execution. Thus, the 100% condition was considered only in the behavioral analysis, whereas neural analyses focused solely on the 50% and 80% predictability conditions.

### 2.3 EEG recording and preprocessing

For EEG recording, a BrainAmp DC 64-channel EEG system with actiCAP active electrodes was used. The sampling rate was 1 kHz, with a 100 Hz online low-pass filter applied. Electrodes were placed according to the international 10/20 system. The FCz channel served as the reference electrode. Two electrodes placed laterally to the eyes’ outer canthi monitored eye movements. During the recording, impedances were kept below 15 kΩ.

EEG data preprocessing was conducted with EEGLAB 21.1. toolbox (Delorme & Makeig, 2004) implemented in MATLAB (2021b, Mathworks, Natick, Massachusetts, USA). Offline band-pass filtering was applied between 0.05 and 80 Hz using a low-pass/high-pass filter cascade of Hamming-windowed sinc filters (EEGLAB pop_eegfiltnew, zero-phase non-causal filter, high-pass filter order = 66000, low-pass filter order = 166). The data were then re-referenced to the average of all EEG channels. The Infomax algorithm for independent component analysis (ICA) with principal component analysis (PCA)-based dimensionality reduction (PCA dimension = 32) was employed for artifact removal (Delorme et al., 2007). ICA components constituting blink artifacts were removed after a visual inspection of their topographical distribution and the frequency contents of the components. A maximum of 3 components per participant was removed in this way (<10%). Malfunctioning channels were identified based on channel spectral characteristics and ICA components. A maximum of 6 malfunctioning EEG channels per participant was allowed without exclusion from the study. These were interpolated using the default spline interpolation algorithm implemented in EEGLAB. Then, the data were low-pass filtered (using a Hamming-windowed sinc filter with a 45 Hz cutoff frequency, a zero-phase, non-causal filter, and a filter order of 294) to attenuate high-amplitude transient noise peaks, and epochs between -500 and +4500 ms were extracted relative to the onset of cue events. Epochs including an amplitude change of 100 µV or greater were rejected from further analysis (ratio of rejected epochs: M=9.7%, SD=10.1%). Because the number of epochs in the 80% predictable condition was higher than in the 50% predictable condition, the number of epochs was equalized between conditions in a within-participant manner. To this end, epochs were randomly removed (Matlab randperm function) from the 80% predictable condition, ensuring the number of remaining epochs was equal to that in the 50% predictable condition for the same participant. The number of epochs varied across participants but was consistent within each condition (M = 69.12; SD = 5.06).

### 2.4 Data analysis

#### 2.4.1 Behavioral data

Statistical analyses were performed in STATISTICA (version 13.1, TIBCO Software Inc., Palo Alto, California, USA). A 2x2 repeated-measures ANOVA was used to compare the effects of cue type (predictive vs. unpredictive) and target type (early vs. late) on reaction time (RT) measured from target onset. Response times longer than 1000 ms from target onset were excluded from the analysis. The alpha level was set at 0.05. Partial eta squared (ηp2) is reported as the measure of effect size. Post-hoc tests were computed by Tukey’s “Honest Significant Difference” (HSD) pairwise comparisons (Tukey, 1949).

#### 2.4.2 ERP cluster analysis

After preprocessing, ERP cluster-based permutation analysis was applied to compare cue- and target-evoked potentials across the two conditions: 50% and 80% predictable.

The cluster-based analysis was performed using the Brainstorm MATLAB toolbox (Tadel et al., 2011; version 11 July 2024). For each comparison, a permutation test was applied to the epochs. In the post-cue period, epochs for the ERP permutation test were defined from 0 to 1300 ms relative to cue onset (i.e., the entire period between cue onset and the subsequent target onset). In the post-target period, epochs for the ERP permutation test were defined between 0 and 1300 ms relative to the onset of early target events to maintain comparability with the post-cue period. The conditions were compared with two-tailed paired t-tests using 1000 permutations. The minimum number of neighboring channels was set to 3, and the alpha level was set to 0.05.

#### 2.4.3 EEG source localization and functional connectivity analysis

The Brainstorm toolbox (Tadel et al., 2011; version 11 July 2024) was used to perform EEG source reconstruction, following the protocol of previous studies (Song et al., 2015; Huang et al., 2016; Pizzagalli, 2007; Tóth et al., 2020). The forward boundary element method (BEM) head model was used as provided by the openMEEG algorithm (Fuchs et al., 1998, 2002; Gramfort et al., 2010). Individual structural MRI and electrode positions were available for each participant and used for head modeling to improve EEG source localization accuracy (Ignatiadis et al., 2022). Anatomical magnetic resonance images (MRIs) were segmented via Freesurfer (Fischl, 2012). The recorded activity was mapped to the cortical surface via dynamic statistical parametric mapping (dSPM) (Dale et al., 2000). The noise covariance was calculated from a 200-ms interval preceding the onset of cue events. Dipole orientations were considered as constrained to the surface, and source signals were reconstructed at 15000 vertices describing the pial surface. By averaging dipole strengths across voxels, the mean source waveforms were obtained for 62 cortical areas (regions of interest, ROIs) based on the anatomical segmentation of the Desikan-Killany-Tourville atlas (DKT atlas, Klein and Tourville, 2012). Mean source waveforms corresponding to the 62 ROIs were used for FC analyses.

For FC analyses, the time window of interest was selected based on the ERP results for the post-target period (see Results): 100-500 ms after target onset. Because the specified window was 400 ms, the reconstructed ROI activities were high-pass filtered at 2.5 Hz to retain the period of the slowest oscillation allowed by the window length. For the post-cue period, the ERP analysis did not yield a significant difference between conditions. Yet, during that period, the anticipatory process is assumed to build gradually and continuously (Walter et al., 1964; Kononowicz & Penney, 2016). Based on this notion, the last 400 ms of the post-cue period (between 900 and 1300 ms after cue onset) was used to maintain comparability with the post-target period. Similarly, the same high-pass filtering (2.5 Hz) was applied as in the post-target period. Finally, epochs were paired by conditions for each participant. The 50%-80% predictable difference waveform was computed for each ROI in each epoch, after pairing epochs randomized separately for each participant (Matlab *randperm* function). For each epoch pair, the waveforms were extracted for each ROI, and the 50% predictable - 80% predictable difference waveform was calculated separately. These different waveforms were used for FC analyses.

FC was analyzed using the Normalized Directed Transfer Entropy (NDTE) framework (Deco et al., 2021; see Ignatiadis et al., 2024, for an application on EEG data). The NDTE framework provides a bidirectional (inflow/outflow) description of the functional information flow. The NDTE flow FXY from time series X to Y (from a source brain area X to another target brain area Y) is defined as the mutual information corresponding to the degree of statistical dependence between the past of X and the future of Y, normalized by the joint mutual information that the past of both signals together, X and Y, have about the future of Y. The normalization enables one to compare and combine the directed mutual information flow across different pairs of brain regions. The mutual information can be calculated from conditional entropies, and the conditional entropies can be estimated from the covariance matrices of X and Y (Brovelli et al., 2015). Following Deco et al. (2021), we estimated the order (or “maximum lag”) of the NDTE model by fitting the autocorrelation function to the first minimum across conditions and participants. The resulting average decay times were 15.09 ± 4.19 ms for the post-cue period and 14.56 ± 2.62 ms for the post-target period. These values determined the integration time window used to estimate directed information flow between regions.

In our study, NDTE values were tested for statistical significance and standardized using circular-shift surrogate data following the method proposed by Deco et al. (2021; 2022); and applied to EEG by Ignatiadis et al. (2024). The advantage of time-shifted surrogates is that the properties (e.g., amplitude and autocorrelation spectrum) of each signal are maintained because the signals are the same, but the cross-correlation between the signals is reduced to chance level (Quiroga et al., 2002). Following the method proposed by Deco et al. (2021), 100 independent circular time-shifted surrogate iterations were performed for each considered ROI pair in each trial. Statistical significance of connections between ROIs was calculated using p-value aggregation via Stouffer’s method (Stouffer et al., 1949) across trials within a participant, and subsequently across all participants. After aggregating p-values across trials and participants, the correction for multiple comparisons was performed using the false discovery rate method of Benjamini and Hochberg (1995). The corrected values were then used to create a binary matrix indicating, with 1s or 0s, whether the corresponding connection was significant. Finally, significant NDTE values were standardized by subtracting the mean and dividing by the standard deviation of the corresponding 100 surrogate NDTE values.

Standardized NDTE values were used to identify functional hub regions in the brain. Following the concept of the FRIC introduced by Deco et al. (2021), we aimed to determine the core set of brain regions that are more functionally densely connected among themselves than to other brain regions. As the first step in the FRIC analysis, standardized NDTE values were averaged across trials and participants, yielding a single standardized NDTE matrix of dimensions ROI x ROI. For each ROI, the total inflow from all ROIs of the cortical parcellation is defined as the sum of connectivity across all columns of the matrix. The total outflow per ROI is defined as the sum of connectivity across all rows of the matrix.

The major hubs were defined through an iterative process. After sorting the regions by inflow from highest to lowest, the largest subset of ROIs was identified that had a significantly larger FRIC-value than any other set with the same number of regions. The FRIC-value is defined as the sum of connections between all club members, plus the sum of all incoming connections of the club members, minus the sum of all outgoing connections of the club members. The significance of the FRIC-value corresponding to a club with k members was assessed via 100000 Monte Carlo simulations for each tested value of k. In each permutation, surrogate clubs with k members were created, each having the same k-1 members and one additional random member. Starting with the ROI with the largest incoming NDTE flow, the algorithm continued to consider larger clubs as long as the p-value of the comparison between the considered club and the surrogate clubs was less than 0.05.

The mathematical formulations of the NDTE framework and the FRIC method, as proposed by Deco et al. (2021), are presented in the Appendix.

The brain networks were visualized with the BrainNet Viewer (https://www.nitrc.org/projects/bnv/, Xia et al., 2013).

#### 2.4.4 Correlating Reaction Time Differences with Functional Connectivity

Reaction time differences between the 50% predictable and 80% predictable conditions were correlated with each of the strongest 20 functional inflow connections (∼0.5 % of all edges) using Pearson’s correlation coefficient. The correlation was calculated for functional inflow connections in both the post-cue and post-target periods.

## 3 Results

### 3.1 Behavioral results

The task design included three target types based on their within-trial predictability: 1) 50% predictable targets: early targets following an unpredictive cue, 2) 80% predictable targets: early targets following a predictive cue, and 3) 100% predictable targets: late targets, which occurred if an early target was absent (Figure 1A and B).

A repeated-measures ANOVA was conducted on RTs with two within-participant factors: CUE TYPE (predictive vs. unpredictive) and TARGET TYPE (early vs. late). The analysis revealed a significant interaction between CUE TYPE and TARGET TYPE, *F*(1,19) = 11.20, *p* < 0.01, η²ₚ = 0.37, as well as a significant main effect of CUE TYPE, *F*(1,19) = 5.92, *p* < 0.05, η²ₚ = 0.24. Post-hoc comparisons confirmed that RTs were significantly slower in the 50% predictable condition compared to the 80% predictable condition, *t*(19) = 4.97, *p* < 0.001, Cohen’s *d* = 0.70 (Figure 1C).

As predicted, RTs were significantly faster for 100% predictable targets (condition 3: late targets) than for 50% predictable targets (condition 1), *t*(19) = 5.05, *p* < 0.001, Cohen’s *d* = 0.58. In contrast, no significant difference was found between 80% and 100% predictable targets (conditions 2 and 3), *p* > 0.05, suggesting that high predictability minimized RT differences in the predictive cue condition.

### 3.2 ERP results

Figure 2A and B illustrate the averaged ERPs elicited by the cue from the channel locations of the frontal and parietal regions (frontal: Fp1, Fp2, AF3, AF4, AF7, AF8, Fz, F1, F2, F3, F4, F5, F6; parietal: Pz, P1, P2, P3, P4, P5, P6, POz, PO3, PO4, PO7, PO9) that contributed most prominently to the significant spatiotemporal clusters shown in Figure 2C.

**Figure 2.**
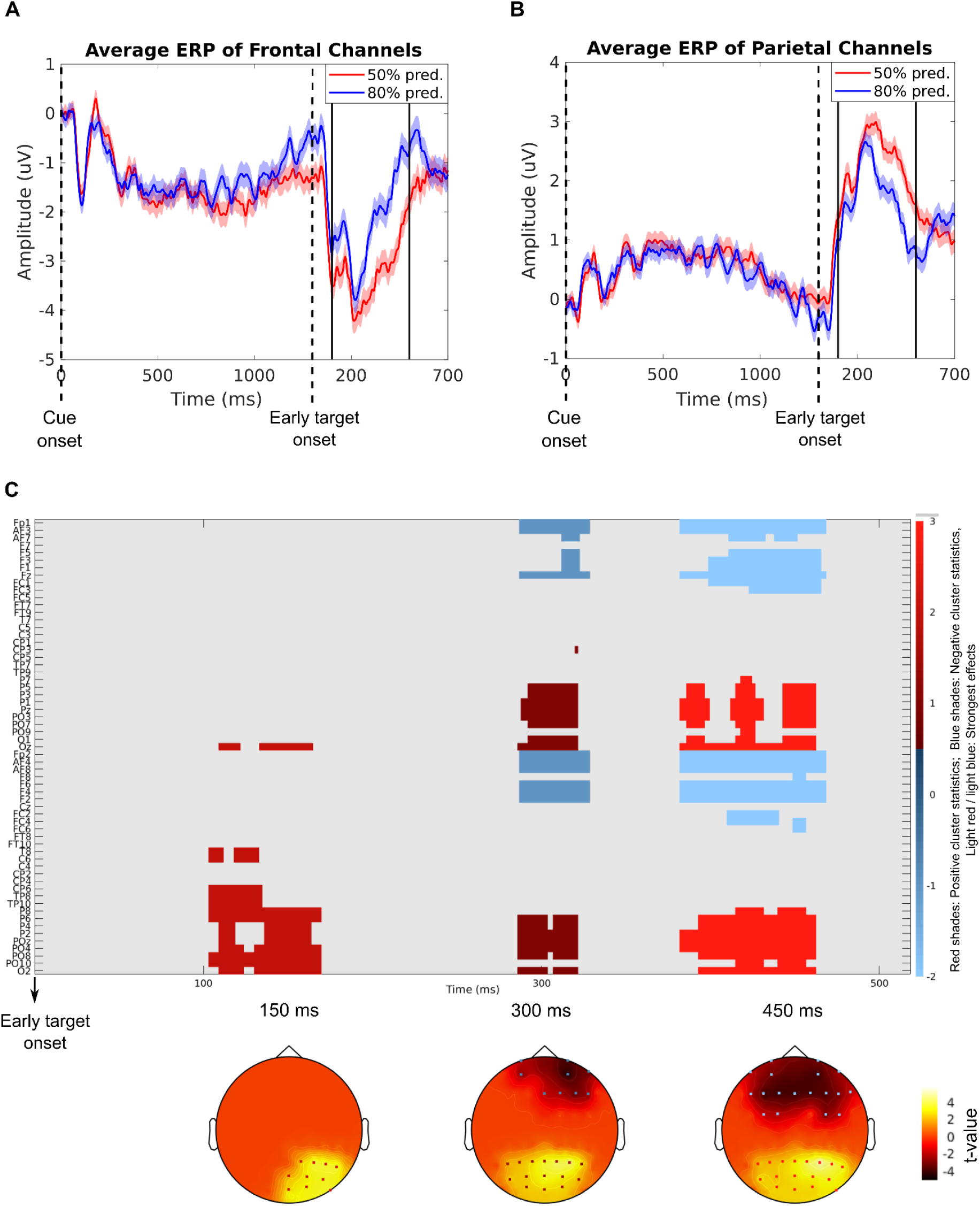
A) cue- and target-evoked ERPs at the average of the frontal electrodes (Fp1, Fp2, AF3, AF4, AF7, AF8, Fz, F1, F2, F3, F4, F5, F6) and B) the averaged parietal electrodes (Pz, P1, P2, P3, P4, P5, P6, POz, PO3, PO4, PO7, PO9) (panel B) showing the most pronounced difference between the 50% predictable (red) and 80% predictable (blue) conditions. Shaded areas denote the standard error. The time range showing the most pronounced difference in the cluster-based permutation test (1400 to 1800 ms after cue onset) is indicated by black vertical lines. C) Results of the cluster-based permutation test on ERPs elicited by 50% predictable vs. 80% predictable conditions (color scale in arbitrary units). Abscissa: Time in seconds (from 1.3 to 1.8 s after cue onset, where 1.3 corresponds to the target onset). The detected clusters span from ∼1.4 s to 1.8 s— ordinate: EEG electrode names. The color scale indicates the direction and strength of the cluster statistics. Red shades: Positive cluster statistics (i.e., 50% predictable > 80% predictable); Blue shades: Negative cluster statistics (i.e., 50% predictable < 80% predictable), Light red/light blue: Strongest effects. The topographical plots show the distribution of significant clusters at 150, 300, and 450 ms.

The cluster-based permutation test comparing the two early target conditions revealed significant differences in the post-target period. In contrast, no significant clusters were observed in the post-cue interval (from cue to target). A prominent post-target cluster was observed between approximately 100 and 500 ms after target onset (i.e., 1400–1800 ms after cue onset), with distinct topographical patterns. Frontal electrodes (e.g., Fp1, Fz, AF3, F1) showed more negative potentials in the 50% predictable condition, while parietal electrodes (e.g., Pz, PO3, PO7, P4) showed more positive potentials for the same condition.

### 3.3 Functional connectivity results

The NDTE framework was used to quantify directional functional connectivity between source-localized cortical regions for the EEG difference between early targets following 80% (predictive) and 50% (non-predictive) cues. Two key time windows were tested: the post-cue period (900–1300 ms after cue onset) and the post-target period (100–500 ms after target onset).

Identified FRIC hubs are shown in Fig. 3, and individual group-level average inflow edges of the identified hub regions showing the highest inflow (the strongest 20 edges [∼0.5 % of all edges]) are illustrated in Fig. 4. These connections are predominantly long-range fronto-temporal, parieto-frontal, and parieto-temporal connections towards FRIC hub regions.

**Figure 3.**
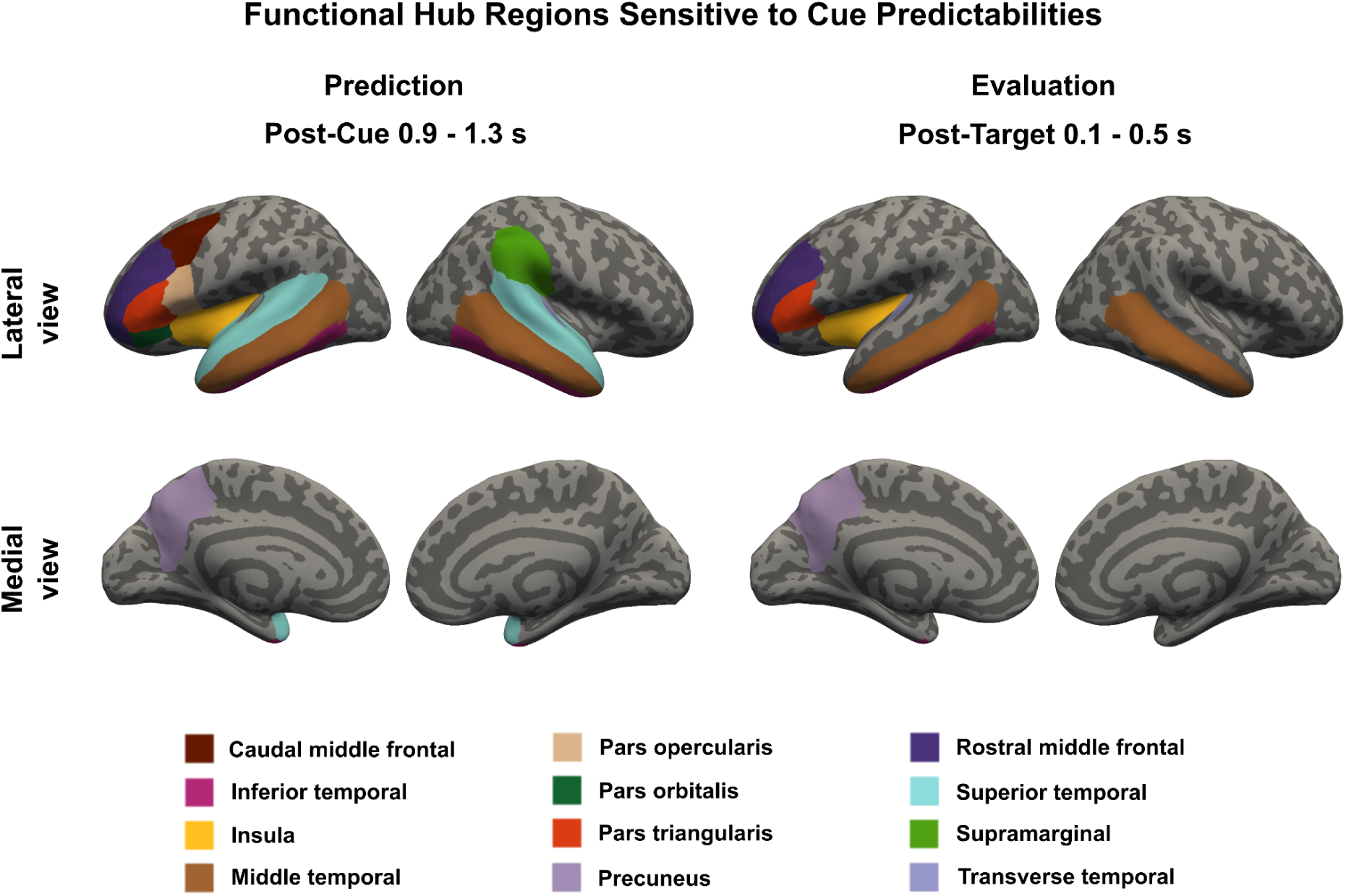
Identified connectivity hub regions for broadband, post-cue 0.9 - 1.3 s and post-target 0.1 - 0.5 s data, based on standardized NDTE values averaged over all epochs of all participants.

**Figure 4.**
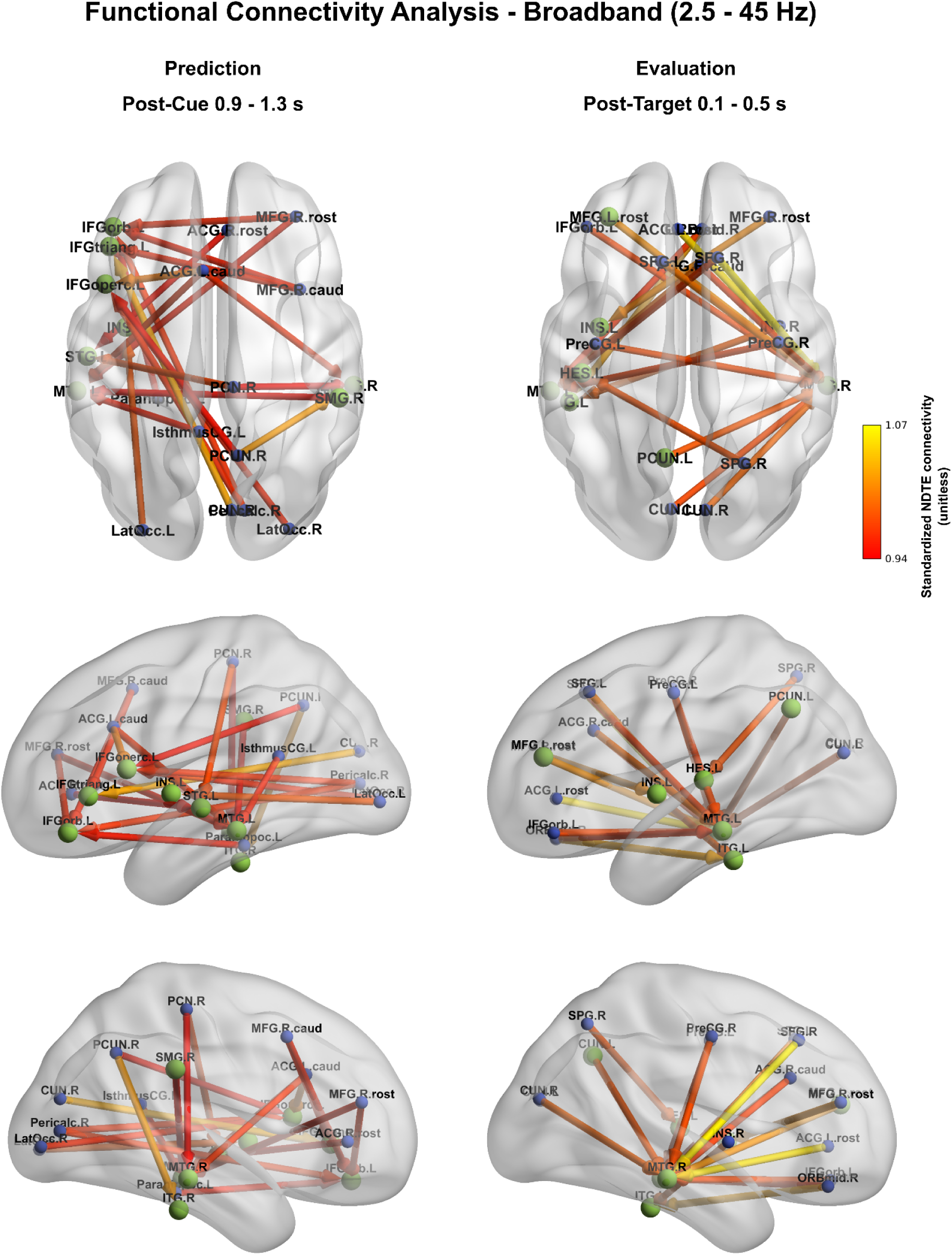
Temporal predictability-related functional inflow connections of different brain regions (group-level average). Regions sensitive to temporal prediction processes after cue or early target stimuli are denoted by green (FRIC hub regions - strongest 20 edges [∼0.5 % of all edges]) and blue colors (non-hub areas). Edge color and edge width denote standardized NDTE connectivity strength (unitless).

During the target prediction window (900–1300 ms after cue onset), FRIC analysis identified a set of predominantly frontal and temporal regions as hubs (Fig. 3, left panel). These included: Frontal - Left rostral middle frontal (rMFG), left caudal middle frontal (cMFG), left pars triangularis, left pars opercularis, left pars orbitalis; Temporal - Bilateral superior, middle, and inferior temporal cortices, right transverse temporal cortex; Other - Left insula, right supramarginal gyrus, and left precuneus. The strongest 20 inflow connections during the prediction window (Fig. 4, left panel) revealed a predominantly left-lateralized network, with dense inputs to frontal and temporal hub regions. These connections are predominantly interhemispheric. Directed edges prominently connected the middle and superior temporal, as well as the left inferior frontal hubs, to parietal, occipital, and middle frontal areas, consistent with anticipatory processing within the dorsal auditory stream under increased temporal predictability.

In the target evaluation window (100–500 ms after target onset), the FRIC network emerged as a set of predominantly temporal and left frontal regions, which served as hubs (Fig. 3, right panel). The hubs included: Frontal - Left rostral middle frontal, left pars triangularis; Temporal - Bilateral middle, and left inferior temporal cortices, left transverse temporal cortex; Other - Left insula, left precuneus. During the post-target window (Fig. 4, right panel), most inflow paths were directed to the left and right middle temporal as well as left inferior and transverse temporal hubs, receiving information from frontal and parietal regions.

### 3.4 Correlation between Reaction Time Difference and Functional Connectivity

The correlation between the 50%-80% predictable condition difference in reaction time and the strongest 20 functional inflow connections was calculated separately for the post-cue and post-target periods (Fig. 5). These functional connections differentially correlate with the observed predictability-related reaction time enhancement. A strong relationship between connectivity and behavioral outcome was observed for fronto-temporal (right rostral middle frontal - left middle temporal, right caudal anterior cingulate - left middle temporal), parieto- and occipito-frontal (right cuneus - left left pars triangularis, right lateral occipital - left pars opercularis), parieto-temporal (right paracentral - left superior temporal) connections and for the left precentral - right middle temporal connection, as well as within the left temporal region (left transverse temporal - left middle temporal). Although the absolute value of these correlation values exceeded 0.4, none of them reached statistical significance after correcting for multiple comparisons; only the correlation of the post-target left transverse temporal - left middle temporal connection with reaction time was found to be marginally significant (*p* = 0.090 after correction, marked by an asterisk in Fig. 5).

**Figure 5.**
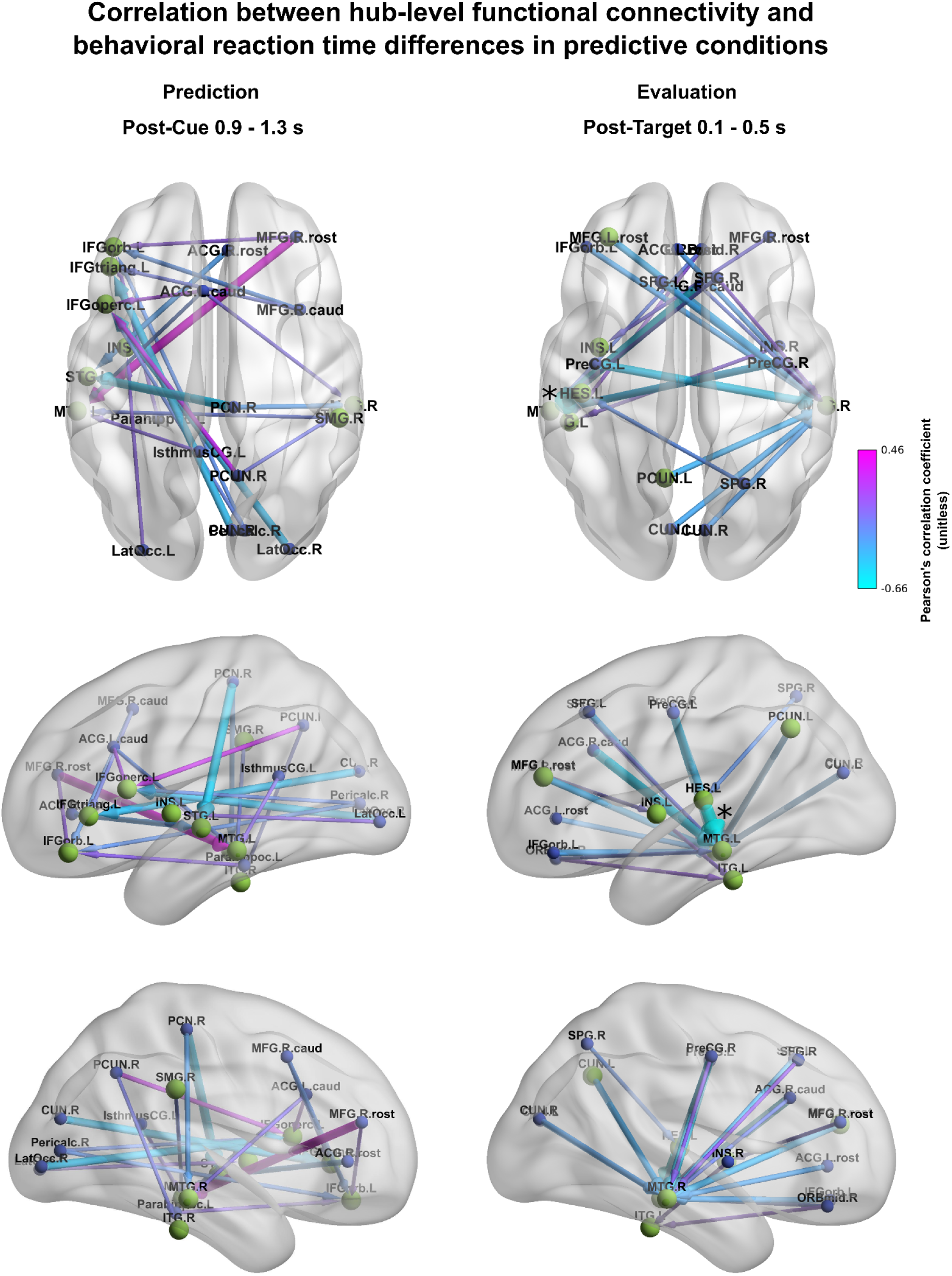
Correlation between the 50% predictable - 80% predictable reaction time differences and the strongest 20 functional inflow connections (∼0.5 % of all edges) in the post-cue (left) and post-target (right) period. Hub regions supporting temporal prediction processes after cue or target events are denoted in green with larger node size (non-hub areas are denoted in blue with smaller node size). Edge color denotes Pearson’s correlation coefficient (unitless), edge width denotes the absolute value of correlation. The marginally significant correlation is denoted by an asterisk.

## 4 Discussion

The present study investigated how large-scale brain networks support temporal prediction and target evaluation during auditory perception by examining the directional functional connectivity on EEG data. By manipulating the predictability of target timing, we identified distinct cortical hubs engaged during prediction versus evaluation, shedding light on the dynamics of predictive processing. Consistent with models of neural entrainment and predictive coding (Schroeder & Lakatos, 2009; Arnal & Giraud, 2012; Benedetto et al., 2021), our results show that temporal prediction engaged a distributed fronto–temporal–parietal network, with prominent hubs in the left inferior and middle frontal gyri, superior temporal cortex, supramarginal gyrus, and insula during the post-cue prediction interval. These hubs, particularly within the frontoparietal control network, showed connectivity patterns that correlated with behavioral facilitation (i.e., faster reaction times), suggesting that temporal expectation enhances sensory readiness via top-down modulation. In contrast, post-target evaluation processing involved greater left-lateralized fronto-temporal connectivity, including the transverse temporal and middle temporal gyri, and was associated with faster responses. This dynamic shift from distributed prediction to targeted evaluation suggests that temporal prediction is not merely a function of local oscillatory entrainment. Instead, it emerges from coordinated activity across multiple large-scale networks that adapt to task demands. Predictive processing may rely on dynamic reorganization driven by prediction signals from frontal anticipation areas rather than solely on distributed sensory updating and association areas.

### 4.1 Predictive timing accelerates behavior

The present findings demonstrate that temporal prediction significantly improves response speed, aligning with theories of perceptual prediction that suggest the brain optimizes processing efficiency when event timing is predictable. Replicating Stefanics et al. (2010), reaction times (RTs) were inversely related to target-onset uncertainty. Early targets preceded by an 80 % predictive cue, and late targets (100 % predictable once the early window elapsed), elicited markedly faster RTs compared to early targets in the 50 % condition. These findings align with behavioral work showing that temporal certainty sharpens perceptual readiness (Bar, 2007; Jones, 2018) and indicate that participants continuously update temporal priors when early events fail to occur (Bubic et al., 2010).

Additionally, the results suggest that participants dynamically updated their temporal expectations. In trials where an early target did not appear, participants shifted their expectation towards the late target, yielding similar RTs for late targets and early targets predicted with high confidence. This finding highlights the brain’s ability to adapt to environmental probabilities and optimize sensory processing based on updated temporal information. Overall, these findings provide strong evidence for the impact of temporal predictive processing on target detection, showing that the brain’s ability to form temporal predictions enhances sensory efficiency and responsiveness. The significant differences in RTs between conditions of varying predictability emphasize the importance of temporal cues in guiding attention and preparing the sensory system for imminent events (Jones, 2018; Large & Jones, 1999).

### 4.2 Event-related dynamics during target detection

The ERP results show that implicit temporal expectancy mainly modulates post-target processing in a distributed fronto-parietal network. A robust cluster from ∼100–500 ms after target onset exhibited more positive parietal activity and more negative frontal activity for 50% relative to 80% predictable targets, whereas no reliable cue-locked differences were detected within the cue–target interval. This pattern suggests that when temporal predictions are weak, later evaluative and decision-related processes require a greater computational load than early sensory gain mechanisms.

Early sensory components (N1, ∼90–130 ms; P2, ∼150–230 ms) typically reflect sensory gain and perceptual consolidation and can be enhanced under explicit temporal preparation (Schroeder & Lakatos, 2009; Polich, 2003, 2007). In our implicit design, however, these deflections showed no reliable modulation, consistent with reports of mixed or absent N1 effects under probabilistic timing in both audition and vision (Rimmele et al., 2011). This supports the view that implicit temporal prediction operates mainly via oscillatory phase alignment rather than large early ERP amplitude shifts. The less predictable intervals between the cue and the target likely reduced phase consistency, yielding diffuse preparatory states. Prior work suggests that low-frequency entrainment may still be functionally relevant even when subthreshold in scalp averages (Lakatos et al., 2008; Stefanics et al., 2010).

By contrast, later evaluative processes were susceptible to predictability. Parietal electrodes exhibited a larger P3b for less predictable targets, consistent with context-updating accounts in which reduced temporal certainty increases model revision (Donchin & Coles, 1988; Polich, 2007). This aligns with auditory and cross-modal findings that low temporal confidence or invalid cues amplify the P3 (Rimmele et al., 2011; Jones et al., 2022). Since target acoustics were identical across conditions, these effects reflect temporal expectancy rather than sensory novelty. Frontal negativity accompanied the parietal P3b, indicating increased executive engagement when predictions were weak. Such fronto-parietal dissociations—greater parietal positivity with frontal control engagement —are typical under temporal uncertainty (Polich, 2007; Rimmele et al., 2011). Importantly, our design manipulated only temporal predictability, extending prior findings beyond joint temporal-spatial manipulations (Rimmele et al., 2011). In sum, implicit temporal expectancy reshaped target processing chiefly at later, decision-related stages. Anticipatory effects were diffuse at the scalp level, whereas parietal updating and frontal control increased under uncertainty. These findings justify moving beyond component-level ERPs to connectivity analyses that can capture distributed anticipatory information flow.

### 4.3 Functional brain networks reflecting the anticipation and detection of auditory events

This study aims to characterize large-scale functional connectivity dynamics that support temporal prediction and subsequent target evaluation using EEG-based FRIC analysis. The results reveal distinct but overlapping sets of cortical hubs engaged during the prediction and evaluative phases. In the post-cue interval, we observed prominent connectivity hubs in the left inferior frontal gyrus (IFG, including pars triangularis, opercularis, and orbitalis), rostral and caudal left middle frontal gyrus (MFG), bilateral superior/middle/inferior temporal cortices, the right transverse temporal cortex (Heschl’s gyrus), the left insula, the right supramarginal gyrus (SMG), and the left precuneus. In contrast, after the target’s appearance, the network of hubs became more left-lateralized, centered on the left rostral MFG and left IFG (pars triangularis), along with bilateral middle temporal gyrus (MTG), the left inferior temporal gyrus (ITG), the left transverse temporal cortex (Heschl’s gyrus), the left insula, and the left precuneus. Notably, we found that the strength of one post-target functional connection – between the left transverse temporal gyrus (primary auditory cortex) and the left MTG – was positively correlated with faster behavioral reaction times, albeit at a marginal level of significance. This suggests that individuals who achieved more effective coupling between early auditory regions and higher-order temporal areas were able to evaluate the target and respond more rapidly, linking stronger post-target connectivity to more efficient performance (facilitative effects of temporal orienting on behavior, Widmann & Schröger, 2022). Taken together, these findings indicate that temporal prediction engages a broad fronto-temporal-parietal network which then evolves into a more focal auditory–frontal circuit for evaluating whether the prediction was fulfilled, a pattern consistent with prior evidence of fronto-temporal interplay in auditory timing and attention tasks (Arnott et al., 2004; Rauschecker & Tian, 2000).

Each identified key cortical hub region during the predictive and evaluation windows is categorized according to its canonical large-scale brain network affiliation (see Table 1).

**Table 1.** Network affiliation of FRIC hub regions identified during the temporal prediction task.

| Observed Region (Hub) | Hemisphere | Network Affiliation | Time Window |
| --- | --- | --- | --- |
| Rostral middle frontal gyrus (rMFG) | Left | Frontoparietal Network | Post-cue, Post-target |
| Caudal middle frontal gyrus (cMFG) | Left | Frontoparietal Network | Post-cue |
| Pars triangularis (IFG, BA45) | Left | Frontoparietal Network | Post-cue, Post-target |
| Pars opercularis (IFG, BA44) | Left | Frontoparietal Network | Post-cue |
| Pars orbitalis (IFG, BA47) | Left | Frontoparietal / Salience Network | Post-cue |
| Superior temporal cortex (STG) | Bilateral | Salience Network | Post-cue |
| Middle temporal cortex (MTG) | Bilateral | Default Mode Network | Post-cue, Post-target |
| Inferior temporal cortex (ITG) | Bilateral, Left | Default Mode Network | Post-cue, Post-target |
| Transverse temporal cortex (Heschl's gyrus) | Right, Left | Salience Network | Post-cue, Post-target |
| Insula | Left | Salience Network | Post-cue, Post-target |
| Supramarginal gyrus (SMG) | Right | Salience / Frontoparietal Network | Post-cue |
| Precuneus | Left | Default Mode Network | Post-cue, Post-target |

**Note.** Network affiliation is based on canonical large-scale functional networks described by Seeley et al. (2007). Abbreviations: MFG – Middle Frontal Gyrus; FPN – Frontoparietal Network; SN – Salience Network; DMN – Default Mode Network; IFG – Inferior Frontal Gyrus; STG – Superior Temporal Gyrus; MTG – Middle Temporal Gyrus; ITG – Inferior Temporal Gyrus; SMG – Supramarginal Gyrus.

Importantly, the set of regions identified here maps onto known large-scale brain networks, providing clues to their functional roles in our task. The left IFG and MFG hubs are core components of the frontoparietal control network, which is recruited by cognitively demanding tasks across domains (Duncan, 2010). Consistent with their roles in executive control and working memory (Seeley et al., 2007; Uddin, 2015), these frontal regions likely served as “flexible hubs” coordinating top-down preparatory processes for temporal prediction (Cole et al., 2013; Marek & Dosenbach, 2018). Such an interpretation aligns with the Flexible Hub theory, which posits that the lateral prefrontal cortex dynamically updates its global connectivity to meet task demands (Seeley et al., 2007; Uddin, 2015). Indeed, recent evidence shows that frontoparietal regions can rapidly adjust their connectivity patterns across task states to guide behavior (Cole et al., 2013; Waskom et al., 2014). In our data, the left lateral prefrontal cortex (IFG/MFG) may serve exactly this function – interfacing with sensory regions to maintain cue-based temporal expectancy and then integrating incoming sensory evidence at target onset. In parallel, the engagement of the left insula (during both anticipatory and evaluation periods), together with the transient involvement of right SMG post-cue, points to recruitment of the salience/ventral attention network (Seeley et al., 2007; Uddin, 2015). The anterior insula is a key node of the salience network that detects behaviorally relevant events and triggers additional resource allocation (Seeley et al., 2007; Uddin, 2015). We speculate that in the predictive interval, the insula (and possibly the right temporoparietal junction region including SMG) monitored for any salient temporal cues or context changes, and at target delivery it helped mediate the switch from an internal predictive state to an external evaluation state (Corbetta & Shulman, 2002; Japee et al., 2015; Vossel et al., 2014). This is consistent with the known role of the ventral frontoparietal network in reorienting attention to unexpected or significant stimuli (Corbetta & Shulman, 2002) – here, the moment of target onset constitutes a critical event that needs to be evaluated, even if it was temporally anticipated. Finally, the consistent presence of the left precuneus (in both phases) and the involvement of lateral temporal cortices (MTG/ITG, especially post-target) may imply the default mode network (DMN) in our task. The precuneus and angular/middle temporal gyrus are established DMN nodes linked to internally generated mentation, autobiographical memory, and contextual associations (Raichle, 2015; Buckner & DiNicola, 2019). Their recruitment suggests that participants engaged brain regions responsible for maintaining an internal model of temporal context and for comparing incoming stimuli against it. In other words, DMN-mediated processes might provide a predictive “context template” against which the awaited target is evaluated. This interpretation dovetails with recent views that DMN activity can support anticipation and scene-modeling even during active tasks (Buckner & DiNicola, 2019); cf. Blank & Davis, 2016. Indeed, the involvement of MTG/ITG could reflect access to learned temporal associations or semantic expectations about the target stimulus, while the precuneus may contribute to mentally projecting the timing interval (a form of “mental time travel”; Barne et al., 2022). When the target occurred, these regions — in concert with the salience and frontoparietal regions — would participate in integrating the sensory evidence with prior expectations, leading to a “knowledge update” once the prediction outcome was clear. Such a mechanism closely parallels the concept of context updating associated with the P300 family of responses (Polich, 2007). In our data, the distributed network connectivity observed post-target (especially the link between auditory cortex and MTG that correlated with faster responses) could be viewed as the neural substrate of updating the mental model: the brain detects that the expected event has occurred (or not) and then rapidly reallocates resources to respond, consistent with P3b-like processes of evaluation and memory encoding (Polich, 2007).

Overall, our findings highlight that temporal prediction and evaluation are supported by the dynamic interactions among multiple large-scale networks rather than by any single region or pathway. This insight can be framed in the context of theoretical models of predictive brain function. On one hand, the frontal-dominant models of predictive coding propose that higher-order frontal regions generate top-down predictions that constrain sensory processing of expected inputs (Friston, 2005; Alamia & VanRullen, 2019). Our observation of prominent left prefrontal hubs during the post-cue interval accords with this view: it suggests that the brain’s predictive state was maintained by the frontal cortex, which likely sent anticipatory biasing signals to posterior regions (e.g., to the auditory cortex). Indeed, evidence from attention research shows that top-down oscillatory activity can be initiated in the frontal cortex and propagate posteriorly to modulate sensory areas (Alamia et al., 2024), providing a mechanistic basis for frontal-driven predictions. On the other hand, our data equally support distributed and hybrid integration models of predictive processing (Kok et al., 2012; Hohwy, 2013; Clark, 2013). These models argue that prediction is an emergent property of interactions across hierarchical levels of the brain rather than a one-way imposition from the top. Figure 4 reinforces this interpretation by showing that the directionality of the strongest functional connections differs between phases: during prediction, most connections converge toward inferior frontal and temporal hubs, whereas during evaluation, the dominant flow shifts outward from frontal regions to temporal cortices. This dissociation in directional flow supports the view that frontal hubs actively generate predictions, while evaluation relies more on downstream updating mechanisms. The involvement of auditory cortices, inferior parietal regions, and DMN hubs alongside the frontal cortex in our task suggests that predictive coding is implemented via a dialogue between multiple regions: frontal areas may initiate and orchestrate predictions, but sensory and associative regions concurrently generate predictions at their own level and send feedback (or prediction errors) to update the higher areas (Clark, 2013; cf. Hohwy, 2013). This cooperative view resonates with frameworks such as “hierarchical predictive coding” and the “Bayesian brain,” in which each level of the cortical hierarchy refines predictions through the reciprocal exchange of signals (Friston, 2005; Kok et al., 2012). In our results, the co-activation of the FPN, SN, and DMN during the task can be seen as evidence for such an integrative approach: the frontoparietal network provides cognitive set-maintenance and error monitoring (Duncan, 2010; Cole et al., 2013), the salience/ventral network injects flexibility by detecting deviations and redirecting attention (Seeley et al., 2007; Vossel et al., 2014), and the default mode/temporal network contributes stored knowledge and contextual guidance (Buckner & DiNicola, 2019). Notably, the coordination among these networks is likely dynamic and time sensitive. Recent studies have demonstrated that attention and prediction involve rhythmic fluctuations in neural coupling, for example, intrinsic alpha/theta-band cycles that periodically gate information flow between frontal and sensory regions (Alamia et al., 2023; Rimmele et al., 2011; Fiebelkorn et al., 2019). It is tempting to speculate that the FRIC-derived connectivity changes we observed might reflect such oscillatory coordination mechanisms, in which the brain periodically synchronizes key regions at opportune moments to optimize processing of the expected target. In line with this idea, the temporal structure of our task (with fixed cue–target intervals) could engage entrainment of neural oscillations (e.g., delta/theta) that involve both higher-order networks and auditory cortex, as shown in prior work on temporal expectations (Rimmele et al., 2011; Jones, 2018). Although our analysis was not frequency-specific, the broad agreement of our findings with these diverse theoretical accounts – from frontal executive prediction to distributed network integration – underscores a hybrid perspective: the brain’s predictive machinery appears to rely on frontal “hub” regions and distributed cortical circuits working in concert to anticipate and evaluate forthcoming events (Nobre & van Ede, 2018; Deco et al., 2021).

From an auditory neuroscience standpoint, our results also enrich the discussion through the lens of the Auditory Event Representation System (AERS) framework. Winkler and Schröger (2015) proposed the extended AERS as an integrative model in which the auditory system builds hierarchical representations of sound sequences and actively predicts upcoming sounds based on both learned regularities and top-down intentions. The present findings support this framework by demonstrating that intentional temporal prediction (driven by an instructive cue) can reconfigure functional connectivity within the auditory-processing network. In particular, the AERS model emphasizes that action intentions can generate expectations for specific sensory outcomes, effectively embedding predictions into the auditory stream. Recent studies have affirmed this notion: for example, Korka et al. (2022) showed that when listeners intentionally produce sounds, the brain treats the intended sound outcome as a prediction that can modify standard deviance-detection responses. In their work, making a sound “predictable” via one’s own action led to an attenuation of the mismatch negativity (MMN) and changes in the P3a, indicating that the violation of an intention-based prediction elicits a different neural response than an externally generated surprise (Widmann & Schröger, 2022). Such findings align with the idea that the auditory system incorporates the “perceptual idea” of a forthcoming sound into its processing (Winkler & Schröger, 2015). Our connectivity results mirror this principle: during the prediction phase, we observed strengthened fronto-temporal links (e.g., between the left IFG and the auditory cortex), which can be interpreted as the neural implementation of an “intended perception” – the brain setting up a template for the expected tone at a specific time. When the target tone actually arrived, connectivity between the primary auditory cortex (left primary auditory cortex) and higher auditory regions (left MTG) was not only prominent but also predictive of faster reaction times across participants. This suggests that those individuals who most effectively aligned early auditory processing with top-down predictions (via strengthened coupling to MTG, a region implicated in contextual and semantic processing of sounds) were better at evaluating the target and initiating a response. In AERS terms, we could say that a well-coupled hierarchy (from Heschl’s gyrus up to lateral temporal cortex and down again) allowed the sensory evidence to be compared to the intention-based prediction more rapidly, resulting in quicker decision and action. This finding, although modest in effect, provides an interesting link between functional connectivity and behavior, echoing the idea that successful prediction can facilitate perception (Kok et al., 2012). That top-down auditory predictions (e.g., generated by one’s own intentions or by explicit cues) can optimize the processing of expected sounds (Jones, 2018; Schroeder & Lakatos, 2009; Arnal & Giraud, 2012; Benedetto et al., 2021). In summary, interpreting our results through the AERS and related predictive coding frameworks suggests that the brain’s response to an anticipated event is not a simple reflex to a cue, but rather a complex preparatory state where frontal, temporal, and insular regions jointly embody an “if–then” model: if the target occurs at the expected time, then here is how to process it. Our empirical evidence of distinct hub configurations for prediction vs. evaluation, and the integration of an intention-based prediction into sensory processing, provides new support for these auditory prediction models (Widmann & Schröger, 2022).

It is important to note that while our use of high-density EEG, realistic head models, and group-level FRIC analysis improves localization reliability (Michel & Brunet, 2019), fine-grained anatomical distinctions—such as between adjacent gyri—should be interpreted cautiously. Additionally, our fixed temporal structure may have induced rhythmic entrainment effects not explicitly modeled here. To refine spatial precision and validate the identified hubs and interactions, multimodal approaches (e.g., EEG–fMRI or MEG) are essential. Simultaneous EEG–fMRI has shown promising correspondence, with studies reporting ∼40% overlap between EEG- and fMRI-derived networks (Abreu et al., 2020). Future research combining these modalities could help clarify whether the observed FRIC hubs—such as in the left IFG or precuneus—align with canonical control and DMN nodes across varying predictive contexts and sensory modalities.

In conclusion, this study provides a comprehensive characterization of the brain’s large-scale network dynamics during the prediction and evaluation of a timed auditory event. We identified how multiple neural networks – including the frontoparietal control network, salience network, and default mode network – jointly orchestrate temporal predictions and interpret incoming sensory information. The left frontal cortex emerged as a significant hub for setting up predictions, in line with frontal-centric theories of predictive coding. Nevertheless, the concurrent engagement of temporal and parietal regions underscores that predictive processing is distributed across the brain’s hierarchy. This integrated perspective bridges theoretical frameworks from the predictive coding literature (Friston, 2005; Clark, 2013) with empirically observed network patterns, suggesting that cognitive brain networks operate in concert to minimize uncertainty about the future. Moreover, our findings extend auditory prediction models (Winkler & Schröger, 2015) by showing that intention-based expectations can modulate inter-regional connectivity and potentially facilitate behavior. While limitations of EEG source localization warrant cautious interpretation of the anatomical specifics, the overall convergence of our results with known functional networks and predictive processing models attests to the value of EEG connectivity mapping in cognitive neuroscience. The present findings advance our understanding of how the human brain proactively prepares for and evaluates events in time, revealing a tightly integrated network architecture that enables us to generate predictions, detect their outcomes, and update our internal models accordingly.

## Data Availability Statement

The datasets analyzed for this study can be found in the OSF repository: https://osf.io/sa7zf/

## Author Contributions

P.N.: Writing – original draft, Writing – review & editing, Software, Data curation, Methodology, Formal Analysis, Visualization; P.K.: Writing – original draft, Writing – review & editing, Data curation, Methodology, Formal Analysis, Visualization; Á.B.: Writing – original draft, Writing – review & editing, Conceptualization, Investigation, Software; O.Sz.: Writing – original draft, Writing – review & editing, Formal Analysis; R.B.: Writing – original draft, Writing – review & editing, Methodology, Supervision; K.I: Writing – original draft, Writing – review & editing, Software, Data curation, Methodology; I.W.: Writing – original draft, Writing – review & editing, Conceptualization, Supervision; B.T:.: Writing – original draft, Writing – review & editing, Conceptualization, Investigation, Methodology, Supervision, Project administration

## Acknowledgments

The authors thank the research assistants Barbara Matulai and Emese Várkonyi for recording the data.

## Appendix

The NDTE flow *F_XY_*from time series *X* to *Y* (from a source brain area to another target brain area) is defined as

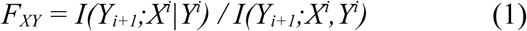

where the mutual information *I(Y_i+1_;X^i^|Y^i^)* corresponds to the degree of statistical dependence between the past of *X* and the future of *Y*, and *I(Y_i+1_;X^i^,Y^i^)* corresponds to the joint mutual information that the past of both signals together, *X* and *Y*, have about the future of *Y*. Normalizing by *I(Yi+1;Xi,Yi)* enables one to compare and combine directed mutual information flows across different pairs of brain regions. With this notation, *Y_i+1_* denotes the activity level of brain area *Y* at time point *i+1*, and *X^i^*denotes the whole activity level of the past of *X* in a time window of length *T* up to and including time point *i* (that is, *X^i^ = [X_i_ X_i-1_ … X_i-(T-1)_]*). Following Deco et al. (2021), we estimated the value of *T* (order, or maximum lag) based on the decay to the first minimum of the autocorrelation function across conditions and participants.

*I(Y_i+1_;X^i^|Y^i^)* can be calculated based on conditional entropies:

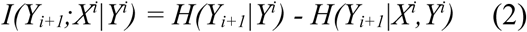

where the conditional entropies are defined as follows:

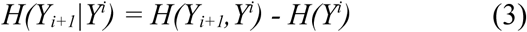

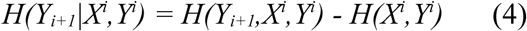

and the conditional entropies can be estimated based on covariance matrices of *X* and *Y* (Brovelli et al., 2015). *I(Y_i+1_;X^i^,Y^i^)* is the sum of the predictability of *Y_i+1_* by the past of *X^i^|Y^i^*and the internal predictability of *Y_i+1_*, that is *I(Y_i+1_;Y^i^)*:

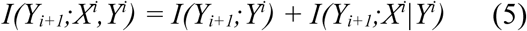

As the first step of the FRIC analysis, standardized NDTE values were averaged across trials and participants, resulting in a single standardized NDTE matrix of dimensions ROI x ROI. Following the notations of Deco et al. (2021), this matrix is denoted by *C_All_*. For each ROI *i* of *C_All_*, the total inflow from all ROIs of the cortical parcellation is defined as the sum of connectivity across all columns of the matrix: *G_in_(i) =* Σ*_j_C_All(i,j)_*. The total outflow per ROI *i* is defined as the sum of connectivity across all rows of the matrix: *G_out_(i) =* Σ*_j_C_All(j,i)_*.

The major hubs were defined through an iterative process. After sorting the regions by inflow from highest to lowest, the largest subset of ROIs *k* was found that had a value *G_FRIC_(k)* significantly larger than any other set with the same number of regions. *G_FRIC_(k)* is defined as follows:

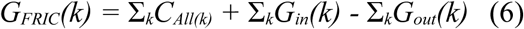

where Σ*_k_C_All(k)_* is the sum of connections between all club members, Σ*_k_G_in_(k)* is the sum of all incoming connections of the club members, and Σ*_k_G_out_(k)* is the sum of all outgoing connections of the club members. The significance value of each *G_FRIC_(k)* was assessed via 100000 Monte Carlo simulations. In each permutation, surrogate clubs with *k* members were created that had the same *k-1* members with one added random new member. Starting with the ROI having the largest incoming NDTE flow, the algorithm continued to confirm FRICs as long as the *p-*value of the comparison between the considered FRIC and surrogate clubs was smaller than 0.05.

